# A complex performance landscape for suction-feeding reveals constraints and adaptations in a population of reef damselfish

**DOI:** 10.1101/239418

**Authors:** Tal Keren, Moshe Kiflawi, Christopher H Martin, Victor China, Ofri Mann, Roi Holzman

## Abstract

The ability to predict how multiple traits interact in determining performance is key to understanding the evolution of complex functional systems. Similar to Simpson’s adaptive landscape, which describes the fitness consequences of varying morphological traits, performance landscapes depict the performance consequences of varying morphological traits. Mapping the population’s location with respect to the topographic features of the landscape could inform us on the selective forces operating on the traits that underlie performance. Here, we used a mechanistic model derived from first principles of hydrodynamics to construct a hypothetical performance landscape for zooplankton prey capture using suction feeding. We then used the landscape to test whether a population of *Chromis viridis,* a coral reef zooplanktivore, is located on a performance peak or ridge based on measurements of kinematic variables recorded *in-situ* during undisturbed foraging. Observed trait combinations in the wild population closely matched regions of high feeding performance in the landscape, however the population was not located on a local performance peak. This sub-optimal performance was not due to constraints stemming from the observed trait correlations. The predominant directions of variation of the phenotypic traits was tangent to the ‘path of steepest ascent’ that points towards the local peak, indicating that the population does not reside on a “performance ridge”. Rather, our analysis suggests that feeding performance is constrained by stabilizing selection, possibly reflecting a balance between selection on feeding performance and mechanical or genetic constraints.

## Introduction

Understanding the mechanisms that shape adaptive diversification is a major goal in evolutionary biology. Because morphology can affect performance, it should respond most strongly to selection on performance (Arnold, 1983; Arnold, 2003; Lande and Arnold, 1983). The earliest framework to assess the fitness consequences of changing phenotype was Simpson’s “adaptive landscape” (Simpson, 1944; Simpson, 1955), which complemented Wright’s adaptive landscape for genotypes (Wright, 1932). This approach consists of mapping all possible phenotypes or genotypes onto fitness, producing a rugged multi-dimensional space in which peaks reflect adaptive combinations surrounded by maladaptive “valleys”. The adaptive landscape was an attempt to conceptualize the complex relationship between multivariate trait combinations and their fitness consequences. In practice, linking phenotypes and fitness has proved difficult, especially when considering multiple traits. As a way to circumvent these difficulties, Arnold (Arnold, 1983) suggested viewing performance as a mediator between morphology and fitness. He further suggested that a performance landscape (i.e. the mapping of all trait combinations to their respective performance values) should provide valuable insights regarding both the selective and non-selective forces involved in trait evolution. This is done through an analysis of the topography of the landscape and the distribution of phenotypic diversity in a population relative to the landscape (Arnold, 2003; Arnold et al., 2001). Conceptually, the performance landscape is similar to the adaptive landscape, with the caveat that performance does not necessarily map linearly to fitness, just as phenotype does not map linearly to fitness. Therefore, the individual performance surface (defined as the derivative of performance with respect to trait value; (Arnold, 2003; Arnold et al., 2001)) is not necessarily equal or proportional to the individual fitness surface (Fear and Price, 1998; Lande and Arnold, 1983)

The topographic features of the performance landscape reflect the selective regimes imposed on performance-determining traits. Stabilizing, disruptive, and correlational selection are reflected by curved landscapes, where performance peaks correspond to local adaptive optima (Arnold, 2003; Arnold et al., 2001). When mapping a natural population onto such a landscape, the distance between the population’s mean and the theoretical peak can provide evidence for constraint, assuming sufficient time for phenotypic evolution (Collar et al., 2009; Hansen, 1997). For example, Arnold and Bennett (Arnold and Bennett, 1988) reconstructed the performance surface that relates crawling performance to body and tail vertebral numbers, and concluded that these two parts of the vertebral column evolve on a performance ridge, defined as an axis on which morphological evolution will result in little change to performance (Arnold, 2003; Arnold and Bennett, 1988). However, generating the landscape by collecting performance data for individuals within the population has a limited ability to infer performance away from the population means, and cannot provide predictions on performance in areas outside the trait space of the observed population. As Arnold put it “It is not clear… that a rugged topography, and the quandries resulting from local peaks, apply in the case of phenotypic characters. More importantly, we have no device that can tell us about landscape features far from the population’s average. Mt. Improbable is, after all, Mt. Unknowable. This is the sense in which the local vision of the landscape is the most tractable.” (Arnold, 2003).

An alternative to reconstructing the landscape is to use mechanistic models of performance. Over the last decades, biomechanical theory and computational methods were used to mechanistically model many aspects of performance such as swimming (Fish and Lauder, 2006; Liao, 2007; Sfakiotakis et al., 1999), running (Barasuol et al., 2013; Kingma et al., 1996; Minetti, 1998), slithering (Hu et al., 2009), flying and gliding (Paranjape et al., 2012; Wu, 2011) among numerous other examples. These models are based on first principles, and produce reliable estimates of performance given a set of phenotypic (often kinematics and morphological) traits that are used as input variables. Because a mechanistic model can (mathematically) solve performance for any input values, it is not necessary to obtain input values from a specific organism, and it is possible to predict performance outside the trait ranges and combinations observed within the population. Furthermore, the data used to generate the landscape is independent of the population, enabling one to generate hypotheses regarding the location of the population relative to the landscape. However, applications of biomechanical models to understand the distribution of performance- determining traits on the performance landscape are rare.

Our goal was to construct and visualize a performance landscape using an empirically-verified mechanistic model of suction-feeding, and map upon it the distribution of individuals within a wild population of *Chromis viridis,* a Coral Reef damselfish which feeds predominantly on zooplankton using suction-feeding.

Mapping the population’s location with respect to the topographic features of the performance landscape could inform us on the selective forces operating on the traits that underlie performance. Specifically, we considered three mutually exclusive scenarios (Fig. 1). Under scenario I, selection for suction performance has a week effect on morphological evolution of the traits that underlie performance, leading to a distribution of phenotypic traits that is unrelated to the performance landscape. In that case, we would expect that the distribution of performance values in the observed population would be similar to that of a population placed at random on the landscape (Fig. 1A). An alternative scenario (II) would be that selection for suction performance has a dominant effect on morphological evolution, and the population sits at the top of a performance peak (Fig. 1B). In that case, we would expect that the distribution of performance values in the observed population will be higher than that of a population placed at random on the landscape, and that the mean individual performance surface is zero. Lastly, under scenario III, the population could be under stabilizing selection, with selection for suction performance countered by trade-offs, genetic of mechanical constraints (Fig. 1B). In that case, we would expect that the distribution of performance values in the observed population will be higher than that of a population placed at random on the landscape and that the selection gradient for each phenotypic trait is different than zero (Fig. 1C).

**Fig 1:**
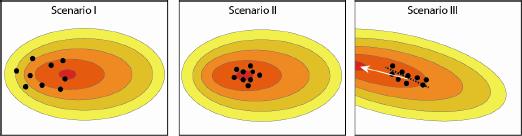
Possible scenarios depicting the distribution of the phenotypic traits with respect to topography of a two-dimensional performance landscape. Trait values for each individual in a population (‘observed’) are depicted as black circles. X and Y axes represent two performance-determining traits, red colors represent higher performance. Under scenario I, selection for suction performance has a weak effect on the traits that underlie performance, leading to a distribution of phenotypic traits that is unrelated to the performance landscape. Under scenario (II), selection for suction performance has a dominant effect on morphological evolution, and the population sits at the top of a performance peak Lastly, under scenario III, the population could be under stabilizing selection, with selection for suction performance countered by trade-offs, genetic of mechanical constraints, or could lay on a performance ridge. In the latter case, the major axis of phenotypic variation (dashed black line) should align with the path of steepest ascent (white arrow), pointing toward the local peak.

## Methods

### Study system

*Chromis viridis* is a common Indo-pacific coral reef species, found at depths of up to 30 m. *C. viridis* live in schools of a few dozen to a few hundred, with adult size ranging from 27 to 59 mm long (Allen and Randall, 1980). Schools are found inhabiting branching corals, feeding on drifting zooplankton in the vicinity of their home coral during the day and sleeping among its branches during the night (Goldshmid et al., 2004).

*Chromis viridis* is an exclusive zooplanktivore, using suction-feeding to capture drifting prey over the coral reefs (Carassou et al., 2008; Coughlin and Strickler, 1990). Prey capture using suction feeding is a widespread and evolutionarily conserved prey capture mechanism among teleost fishes (Day et al., 2015; Wainwright et al., 2015).

Suction feeding can be treated as a hydrodynamic interaction between a solid particle (the prey) and unsteady flows (the suction flow), and can be modeled by tracking the forces exerted on the prey and its trajectory during the strike (Day et al., 2015; Holzman et al., 2012). This hydrodynamic modeling allows us to quantitatively estimate the performance consequences of any given trait combination and to reconstruct a performance landscape independent of the observed population for any phenotype (i.e. any combination of values for the input traits).

The mechanism of suction feeding consists of a swift opening of the mouth and expansion of the oral cavity, which generates a flow of water into the predator’s mouth. Prey that cannot withstand the force of the flow are sucked into the fish’s mouth with the surrounding water (Day et al., 2015; Holzman et al., 2012). Observations, experiments, and hydrodynamic modeling reveal that feeding performance is determined by multiple morphological and biomechanical traits. These traits include the size of the mouth, the speed of mouth opening, and the speed of closing the distance between the predator and prey through forward swimming, and jaw protrusion (Day et al., 2015; Holzman et al., 2008a; Holzman et al., 2012). Many zooplankton species evade those predation attempts by sensing the hydrodynamic disturbance caused by the approaching predator and the suction flows (Buskey et al., 2002; Fields and Yen, 1997; Heuch et al., 2007). In copepods, rapid swimming away from the predator is triggered by the deflection of sensory setae on the antennae in response to the hydrodynamic disturbance. Highly sensitive prey will react to a hydrodynamic disturbance when the predator is still too far to capture it, and could escape (Kiorboe and Visser, 1999; Visser, 2001). Across zooplankton species, sensitivity thresholds differ between species and developmental stages (Green et al., 2003; Kiorboe and Visser, 1999; Visser, 2001). Thus, high feeding performance, from the predator’s standpoint, is its ability to capture sensitive prey by minimizing hydrodynamic disturbance (Gemmell et al., 2014; Holzman and Wainwright, 2009; Viitasalo et al., 1998), which in turn, increases the range of available prey.

### Data acquisition

We filmed prey-acquisition strikes (hereafter “strikes”) of *C. viridis* feeding on naturally occurring drifting zooplankton in their natural habitat. All video sequences were recorded over a period of three months, from November 2013 to January 2014. Videos were recorded during daytime, using natural light, as these fish are visual predators and are only active during the day. Filming was conducted using synchronized high-speed cameras in a waterproof housing (500 frames per second at 1.3 megapixels; Hispec1, Fastec Imaging inc. San-Diego). The underwater video system (Fig. 2) was moved between five schools, each consisting of tens to hundreds of individuals. The cameras were positioned at an angle of ~30° with respect to one another, and were located ~1 m from the coral inhabited by the focal fish school. The two cameras were aimed at the same point in space, ensuring multiple views on objects in the visualized volume. Cameras were calibrated using DLTdv5 (Hedrick, 2008), providing a three-dimensional reconstruction of feeding strikes and enabling time-resolved measurements of distances and locations in a volume of ~20×25×10 cm. The system was re-calibrated every filming session, and measurement error (< 1%) was estimated by filming a ruler that was moved within the visualized volume. Live video feed from the camera was viewed onshore through a tethered cable. A trained observer manually triggered the system to record voluntary prey-acquisition strikes of the naturally foraging fish each time such a strike was observed.

**Figure 2:**
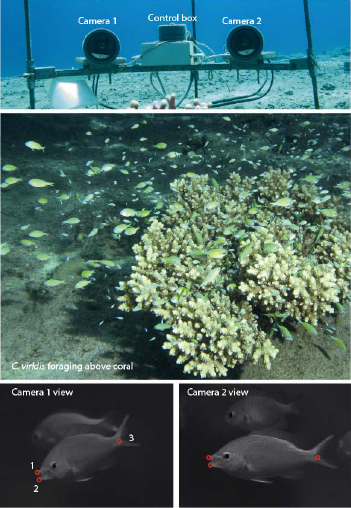
the underwater filming system at the Coral Reef in Eilat (upper panel), was set up to film the zooplanktivorous *chromis viridis* during natural foraging (middle panel). The system captured stereoscopic views of the feeding fish, observed with the two cameras (lower panels). Landmarks (red circles 1-3 in lower panels) were digitized on the fish’s body and head to estimate strike kinematics at 500 frames per second.

For each recorded strike, we digitized four landmarks on the body of the fish: the tip of the upper jaw, the tip of the lower jaw and the base of the pectoral fin (Fig 2, Fig 3). From these landmarks, we calculated the size of the mouth, the location of the mouth center and the location of the fish’s body at each frame. From these measurements we then derived the values of six kinematic traits that are known to be important in determining suction feeding performance: *I*) max gape, defined as the maximal diameter of the fish’s mouth; *II*) time to peak gape, defined as the time it took the fish to open its mouth from 20% to 95% of peak gape; *III*) maximal jaw protrusion measured during the strike, at the fish’s frame of reference; *IV*) time to peak jaw protrusion, defined as the time it took the fish to protrude its jaws from 20% to 95% of maximal protrusion distance; *V*) ram speed, defined as the fish’s swimming speed during the strike; and *VI*) timing of peak protrusion, defined as the time difference between the time of maximal gape and time of maximal jaw protrusion (Fig 3A). Calculations were made as described in (Holzman et al., 2008b). Overall, we analyzed a total of 110 strikes in which we could clearly identify and digitize the landmarks.

The high ratio (~1:20) between the number of fish and the number of sequences recorded in each school translated to a low probability of repeatedly filming the same individual. To further verify data independence, we digitized for each fish the base of the caudal, dorsal, and pelvic fins and used these data to estimate the length and body depth of each individual. Using the measurement error obtained by repeatedly measuring the same fish in several points in space, we calculated the confidence intervals for the measurements of the length and body depth (± 2mm). We then assessed the overlap in size (length and body depth) between all the fish observed in our recordings. We assumed that when an individual is recorded twice, the measures of its length and body depth in the one movie would be within the confidence intervals of these measurements in the other movie. This analysis is conservative, as there could be multiple individuals with the same length and body depth. Overall, < 10% of the strikes in each school were performed by individuals of the same size. Therefore, we regarded the strikes as true replicates.

### Using SIFF to estimate feeding performance

Estimating feeding performance from each set of six phenotypic traits was done using the SIFF model (Holzman et al., 2012). SIFF uses a set of parameters (Fig. 3B) that characterize the prey and describe the kinematics of the mouth and flow speed during the strike to predict the motion of the prey relative to the mouth during suction feeding (Holzman et al., 2007; Holzman et al., 2012; Wainwright and Day, 2007). In this study, we varied SIFF input parameter values based on measured morphology and strike kinematics of *C. viridis* to estimate feeding performance in 3291 phenotypic combinations of the 6 aforementioned traits. These combinations were generated by sampling from the range of phenotypic traits observed in the population, allowing us to construct the performance landscape independently of the observed strikes. As such, we were unconstrained by the distribution or interdependence between traits manifested in the observed strikes. Prey capture performance was defined based on the sensitivity level of the simulated prey that enabled its escape; trait combinations that resulted in capture of highly sensitive prey were thus defined as “high performance strikes” (see below; Fig. 3B,C; (Holzman et al., 2007; Holzman et al., 2012; Wainwright and Day, 2007)).

**Figure 3:**
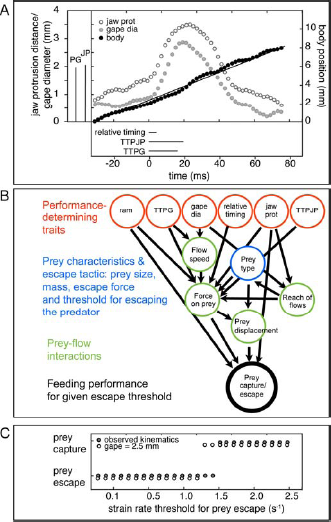
using SIFF to estimate prey capture performance from field-observed and simulated kinematics. Using an underwater high-speed 3D camera system (Fig 2), we recorded body and mouth kinematics of *Chromis viridis* (A). From these, we calculated a set of 6 functionally-relevant traits (red circles in B; values for the presented strike are indicated in boxes to the left and bottom of main panel in A). These traits, in addition to prey’s traits (blue circles in B) were used as input variables for SIFF, a hydrodynamic model of prey capture in fish. The model calculates the displacement of the prey in space and time, based on the forces exerted on it by the suction flows (generated by the fish) and the prey’s propulsive force. For each set of input parameters, the outcome (prey escape/capture) is deterministic. To estimate performance, the model was ran iteratively for each set of input parameters while increasing the strain rate threshold for prey escape (C). At a sufficiently high threshold (causing the prey to escape later in time) the prey is captured, and that threshold is recorded. This process can be carried with observed (filled points in C; a threshold of 1.5 s^−1^) or simulated kinematics, for example using a larger gape diameter (empty circles in C; yielding a threshold of 1.25 s^−1^). To reconstruct the performance landscape, we used ~3300 trait sets, generated computationally by sampling in random from range of observed trait values in the field population.

SIFF is described in detail in (Holzman et al., 2007; Holzman et al., 2012; Wainwright and Day, 2007). In brief, SIFF uses the flow field realized during the strike and a set of parameters that characterize the prey, the body, and the mouth to predict the motion of prey relative to the mouth during suction-feeding, given the suction flow generated by the fish and the ability of the prey to move away from it (Fig 3B). According to SIFF, the movement of the prey relative to the predator’s mouth determines whether prey is captured or not. The total force exerted on the prey is the sum of five component forces: Drag, acceleration reaction force, the force resulting from the pressure gradient across the prey, prey swimming forces, and gravitational forces [the latter will be ignored in the current discussion as most aquatic organisms are approximately neutrally buoyant]. These forces result from the differential in speeds and accelerations between the prey and the water around it, as well as from the gradient of flow across the prey (Holzman et al., 2007; Holzman et al., 2012; Wainwright and Day, 2007).

Flow visualization studies (Day et al., 2015; Holzman et al., 2008b; Staab et al., 2012) reveal that the flow field in front of the mouth can be estimated based on mouth kinematics. We used the linear speed of mouth opening *Gs* (ms^−1^; the derivative of gape with respect to time during mouth opening) to estimate peak flow speed at the center of the mouth *Fs* (ms^−1^) as follows:

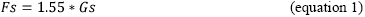

based on the relationships observed in (Holzman et al., 2008b). The temporal pattern of the flow was assumed to follow the gape cycle, with flow starting at 20% and peaking at 95% of maximal gape (Day et al., 2015; Higham et al., 2006; Holzman et al., 2008b). Based on these flow visualization studies, flow speed at any given point in space can be estimated given the flow speed at the mouth center and the instantaneous gape diameter (Day et al., 2015; Higham et al., 2006; Holzman et al., 2008b). From this data, we could derive temporal and spatial flow gradients, and calculate the force exerted on the prey.

A widespread predatory strategy across teleosts to rapidly reduce the distance to the prey is to advance the mouth towards the prey, realized by swimming (ram) and jaw protrusion (Longo et al., 2016). In the context of feeding on evasive zoooplanktonic prey, swimming incurs a cost, because the advancing body produces a hydrodynamic disturbance (sometimes referred to as “bow wave”). Similarly to the suction flows, the bow wave can be sensed by the prey to trigger an escape response (Fields and Yen, 1997; Stewart et al., 2013; Yen et al., 1992). Hydrodynamic disturbance was quantified as strain rate, which can be understood as the rate of deformation of the flow with respect to time. Strain rate is a useful metric for hydrodynamic disturbance because higher strain rates cause faster deflection of sensory antennae of copepods (and other marine invertebrates), ultimately triggering their escape response (Fields and Yen, 1997; Yen et al., 1992). We used observed movements of the body and mouth center as SIFF inputs for the distance between the prey and predator and the properties of the bow wave. The relationship between swimming speed and the hydrodynamic disturbance generated by the body was estimated by visualizing the flow field in front of a formalin-fixed *C. viridis* specimen (standard length = 50 mm) held at different flow speeds (2.5 − 20 cm s^−1^; n= 10 speeds) in a recirculating flume. Flow visualization and analysis followed the protocol described in (Holzman and Wainwright, 2009), except that here the fish was mounted in the flume and not free swimming. These measurements showed that strain rate increased with increasing swimming speed and decreased with the distance to the prey, following the equation:

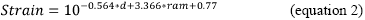

where *strain* is the strain rate (s^−1^) at a distance *d* (mm) from the mouth, and ram is the swimming speed (m s^−1^).

Prey was modeled as a naturally buoyant prolate spheroid (0.1 mm in peak diameter, 2 mm length) located on the centerline across from the mouth. The initial location of the prey at the onset of simulation was determined as the point in space where the mouth center would be at the time of peak gape. The prey was modeled to escape directly away from the predator. Peak escape force (2×10^−5^ N), escape duration (10 ms) and latency (1 ms) were adopted from (Buskey and Hartline, 2003; Buskey et al., 2002). The prey was modeled to escape once it sensed a hydrodynamic disturbance that exceeded a given strain rate threshold. Strain at the location of the prey was calculated as the sum of strain rates resulting from the advancing body and the suction flows (i.e we did not account for a possible interaction between the two sources of flow).

For each set of parameters, SIFF determines whether the prey was captured or escaped. To estimate feeding performance, we iteratively ran SIFF for each set of parameters starting with an unrealistically sensitive prey, which escaped when strain rate exceeded 10^−2^ s^−1^. This threshold ensured that all prey will escape in the first iteration. We then increased strain rate threshold by 5% (thus decreasing prey sensitivity) and re-ran SIFF until the prey was captured, and recorded the strain rate threshold in the last simulation (Fig 3C). This value was used as an estimate of the most sensitive prey that can be captured under a given set of traits, thus representing feeding performance.

A trait set of 3291 phenotypic combinations was generated by randomly sampling from the range of phenotypic traits observed in the population (under a uniform probability distribution). Each trait was sampled independently of the other 5 traits, thus producing sets of combinations that were not observed in the population studied. This sampling ensured dense uniform coverage of the phenotypic space, regardless of the distance from the population mean (Fig S1). Thus, our theoretical calculation of the performance landscape is independent of the trait covariance relationships in the focal *Chromis* population.

### Estimating the contours of the performance landscape

Phenotypic data and performance estimates of this simulated population were used to construct a continuous performance landscape, statistically modeling the relationship between phenotypic traits and performance in the complex, multivariate, mechanism of suction-feeding. To estimate the topography of the performance landscape we used multivariable generalized additive model (GAM) with penalized cubic regression splines, and shrinkage smoothing to further penalize highly smoothed terms (Wood et al., 2015). Thus, shrinkage smoothing reduces the estimated effect of parameters with little explanatory power. Cubic regression splines were chosen over thin plate regression and simple multiple linear regressions as they provided the best fit in terms of deviance explained and prediction of SIFF results.

We used GAM to fit splines that represent, in the multivariate space, the linkage between the six kinematic traits across 3291 simulated individuals with their feeding performance. Given a model structure specified by a GAM model formula, the GAM function attempts to find the appropriate smoothness for each spline term (Wood et al., 2015). The fitted model had an adjusted R-square of 0.899, and explained 90.4% of the deviance. Of the six traits and their paired interactions, only two interactions (maximum jaw protrusion with time to peak gape and with gape-protrusion time difference) were non-significant (and thus were assigned a nearly zero coefficient).

To further test the predictive ability of the GAM surface, we divided the dataset into training/test datasets, containing randomly sampled 3181 trait sets used to reconstruct the landscape (training set) and 110 randomly-sampled trait sets used to test it (test set). We then calculated, for the test set only, a linear regression of strain rate thresholds derived from SIFF vs. those derived from the landscape. The process was repeated 100 times. The mean (± 95% confidence interval) intercept and slope of that regression were −0.004 (±0.33) and 1.009 (± 0.015), respectively, and the R^2^ was 0.88 (± 0.05). Given that the 110 trait combinations of the test set were not used to compile the performance landscape, we concluded that this model is suitable to predict feeding performance for any given phenotypic combination within the observed phenotypic ranges.

### Scenario I: the distribution of phenotypic traits is unrelated to the performance landscape

To test this scenario, we tested whether the feeding performance of the observed population is significantly higher than that expected by chance. We generated a set of simulated populations, by allowing 110 simulated individuals to evolve each trait separately under a Brownian motion model using the observed mean as the starting point. Simulation of Brownian motion was conducted using the package “Geiger” in R statistics (Harmon et al., 2008) using a star phylogeny with branch lengths of 1, and the Brownian rate parameter estimated for the observed population. Thus, this simulation (hereafter “uncorrelated BM”) assumed no trait correlation and no phylogenetic structure within our population. We generated 1000 simulated populations, each with 110 trait sets (simulated “individuals”), and calculated their feeding performance by projecting their trait values on the performance landscape. If the distribution of phenotypic traits is unrelated to the performance landscape, we would expect the median performance of the observed population to fall within the 95% confidence interval of the median performance of the simulated population.

### Scenario II: does the population sit at the top of an adaptive peak?

If the observed median feeding performance is significantly higher than that expected based on the simulated populations, it would be consistent with selection to maximize suction-feeding performance. If the population is located on the performance peak, then a small change in the traits that determine performance should result in a decrease (or no increase) in performance. We mapped the 110 observed sets of traits onto the performance landscape, and estimated the feeding performance for each set. We then calculated the “individual performance surface” following (Arnold, 1983; Arnold, 2003). These performance surfaces are the slope of the performance surface at the individual’s observed location in performance space. Slopes were calculating numerically by adding a small value (0.1 standard deviation of the mean) to one of the six traits, and calculating the difference in performance between the original and shifted set. The process was repeated to calculate the slope for each trait. We further used response surface modeling in the RSM package in R (Lenth, 2009) to locate the stationary point of the response surface, which indicated a performance peak (or trough, depending on the mapped performance; (Brodie et al., 1995)).

### Scenario III: high-performance population resides off the performance peak

If the mean individual performance surface are greater than zero, it would mean that the population is located off the performance peak. In this case, trait variance and covariance could be indicative of the processes generating the off-peak location. For example, trait correlations could limit the realized distribution of performance-determining traits to off-peak locations. To test this hypothesis, we randomized the order of values in each trait, thereby breaking the correlations between this trait and all the other traits. We then compared the distribution of performance (strain rate threshold) to that in the observed population. This was done sequentially for each trait. Additionally, we generated a second set of simulated populations by allowing 110 simulated individuals to move in a correlated Brownian motion manner (using the package “Geiger” in R (Harmon et al., 2008), using the observed mean as the starting point and the observed variance-covariance matrix of the observed population, again assuming no phylogenetic structure within our population (hereafter “correlated BM)”. We generated 1000 simulated populations, each with 110 trait sets (simulated “individuals”), and calculated their feeding performance by projecting their trait values on the performance landscape.

Trait correlations does not necessarily constrain performance. Arnold (Arnold, 2003) suggested that phenotypic traits will evolve along performance ridges that transverse the multidimensional space. If such a ridge in the landscape guides the distribution of the traits in the population, we would expect that the major axis of trait variation in the population to align with the direction of the ridge in the performance space, i.e. parallel (180° or 0°) to the direction to the peak. We therefore used a Response-Surface Method (RSM) (Lenth, 2009) to detect the path of steepest descent. This path is the vector, in the multi-dimension space, which will result in the largest performance increase for a 0.1 standard deviations change in the phenotypic traits (results were identical for 0.5 and 1 SD). We quantified the primary axis of variation in each phenotypic trait using the eigenvectors of PC1 from a correlation based PCA analysis of the 110 observed strikes. We then calculated the angle in the 6- dimentional space, between the two vectors as:

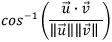

where 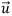 is the vector of coefficients (eigenvectors) for PC1 and 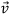 is the vector of steepest descents from the RSM analysis, the numerator is the dot product of the two vectors, and the denominator is the product of the vectors’ norm.

## Results

### Performance and phenotypic attributes of the observed population

Overall, the morphological and kinematic characteristics of suction-feeding in the observed population of *C. viridis* could be described as a swift opening of the mouth (median time to peak gape = 20 ms), to a relatively small maximal diameter (median = 2.6 mm), with a fast jaw protrusion (median time to peak protrusion = 23 ms), reaching its maximal distance (median = 2 mm) 2 milliseconds after the time of max gape, while maintaining a ram speed of ~2 body length per second (median = 77 mm s^−1^). With the exception of maximal jaw protrusion distance, all phenotypic traits were skewed and significantly differed from normal (Shapiro-Wilk’s p<0.05; Supplementary Table S1). Several significant correlations were observed between traits (Spearman rho r > |0.2|; Table 1), including a correlation between maximal gape diameter and max jaw protrusion (r=0.59, p<0.001), between time to peak gape and time to peak jaw protrusion (r=0.73, p<0.001), between time to peak gape and max jaw protrusion (r=-0.4, p<0.001), and between time to peak jaw protrusion and max jaw protrusion (r=-0.37, p<0.001).

**Table 1:**
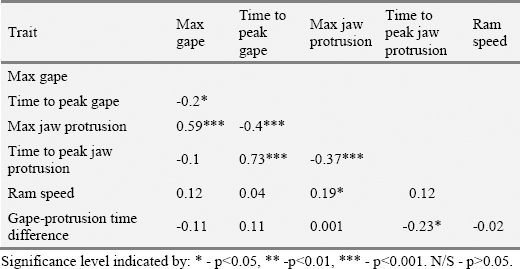
The correlation structure between phenotypic and kinematic traits. With the exception of maximal jaw protrusion distance, all phenotypic traits were skewed and significantly differed from normal (Shapiro-Wilk’s p<0.05; Supplementary Table S1). Therefore, we used Spearman rho for correlation between the traits in the observed population.

The constructed performance landscape revealed a complex, non-monotonic, performance surface (Figure 4; Figure S2) featuring ridges, peaks and flat projections. The density of the contours was different for the different trait combinations, indicating variable performance gradients for each set of traits. Of the six traits and their paired interactions, all but two interactions were significant in the GAM model (Table S2), indicating the importance of trait combinations in determining performance. The range of strain rate thresholds estimated across the landscape is ecologically plausible, ranging from 1-10 s^−1^ (Green et al., 2003; Kiorboe and Visser, 1999; Viitasalo et al., 1998; Visser, 2001), reinforcing the validity of this model in predicting feeding performance based on phenotypic data.

**Figure 4:**
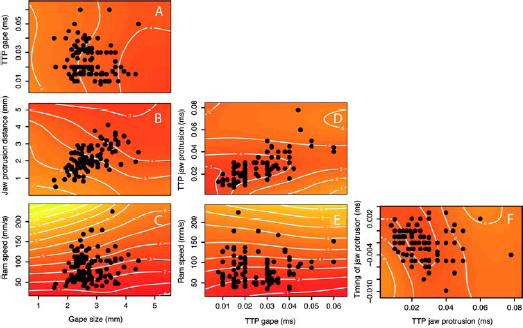
2D projections of the multidimensional performance landscape, generated using a hydrodynamic model of predator-prey interactions. The multidimensional performance landscape was generated based on 3300 simulated prey-acquisition strikes, featuring random trait combinations selected from the observed trait range. Colors (and white contours) represent feeding performance, estimated as strain rate threshold (s^−1^) of the most sensitive prey that can be captured by a predator using that trait combination. Red colors represent high performance (low strain rate thresholds), yellow represents low performance (high strain rate thresholds). Observed trait combinations (black circles, n=110) are overlaid on the performance surface. Note that the 2D visualizations only represent the performance of the featured variables at median values of the other four phenotypic traits, and do not represent trait distribution in the multidimensional space. The plot features six of the 15 possible 2D projections (See Fig S2 for all projections).

### Scenario I: the distribution of phenotypic traits is unrelated to the performance landscape

The median strain rate threshold of the observed population (2.16 s^−1^) was significantly lower by ~25% than the mean median of the 1000 “uncorrelated BM” simulated populations (mean = 2.86; p < 0.001; 95% confidence intervals = 2.58 - 3.09 s^−1^), and was outside the distribution of median performance in the simulated populations (p<0.001; Figure 5A), refuting the hypothesis that the distribution of phenotypic traits is unrelated to the performance landscape.

**Figure 5:**
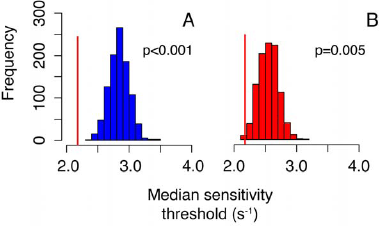
Distribution of performance (strain rate thresholds) in 1000 populations, each with 110 simulated individuals, *versus* performance in the observed population A) Median performance in the observed population (vertical red line) and its distribution in populations generated by an uncorrelated Brownian motion process (“uncorrelated BM” populations; blue bars). B) Median performance in the observed population (vertical red line) and its distribution in populations generated by a correlated Brownian motion process (“correlated BM” populations; red bars). Sensitivity threshold median of the observed population is outside the 95% confidence interval for both simulated populations (p<0.001 for both; n=1000).

### Scenario II: the population sits at the top of an ‘adaptive peak’

While the performance of the observed population was higher than that of the simulated population, the observed population was located off (local and global) performance peaks. This was evident by the individual performance surface calculated for the six traits (Fig 6). Five of the selection surfaces were significantly larger than zero, indicating that a change equivalent to 0.1 sd of the trait value would significantly increase mean performance (decrease threshold strain rate). TTPJP was the only trait for which individual performance surface was not significantly different than 0 (Fig 6D). The RSM analysis identified a local performance peak at peak gape diameter of 5.1 mm, TTPG of 0.019 s, jaw protrusion distance of 6.6 mm, TTPJP of 0.007s (TTPG −0.012), ram speed of 61.8 mm/s and a timing differential of 0.012s. None of the individuals in the observed population had such a combination, however they are within the range observed for similar-sized fish.

**Fig 6:**
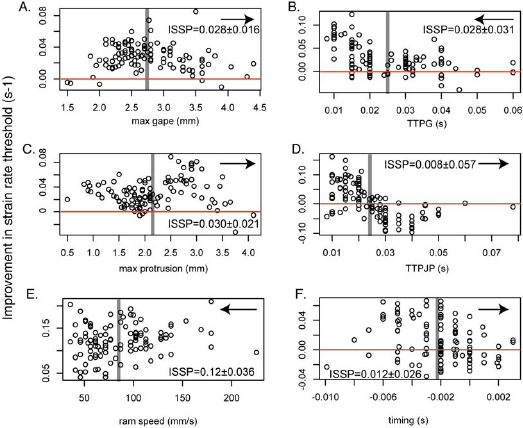
Individual performance surface (Arnold, 2003), as a function of trait values. These selection surfaces represent the slope of the decrease in strain rate threshold (increased performance) following shift of 0.1 standard deviations in trait value. The direction of the change in each trait is indicated by the arrow in the top right corner of each panel. The length of the arrow is proportional to one standard deviations of the trait. Mean (±sd) individual performance surface (ISSP) is depicted for each panel. Vertical gray line depicts the trait mean. The horizontal red line represents ISSP=0; negative values on the y-axis indicate that performance decreased (strain rate threshold increased).

### Scenario III: the population evolves toward the ‘adaptive peak’, but has not reached there

We calculated the angle in the 6-dimentional space between the eigenvector of PC1 of the observed data (explaining ~35% of the total variance), and the vector of the steepest descents, which depicts the direction of change, on the landscape, that will result in maximal performance increase. Contrary to the “performance ridge” hypothesis, that angle was 103.7°, indicating that the major axis of trait variation is tangent to the direction to the peak. A similar result (an angle of 84.5°) was obtained when we calculated the angle between the vector of steepest descents and the vector sum of the coeficients for PC 1-2.

### Are trait correlations responsible for the off-peak location of the population?

Trait correlations can limit the distribution of traits in the trait space and therefor constrain performance, providing a possible mechanism for the off-peak location of our population. However, in contrast to this expectation, observed trait correlations contributed to feeding performance. Simulating populations with the observed trait correlations resulted in a higher performance (lower strain rate threshold) compared to the “uncorrelated BM” populations. However, the median strain rate threshold of the “correlated BM” populations was still significantly higher by ~15% than the observed population (mean 2.55 s^−1^; 95% confidence intervals for = 2.30 - 2.81 s^−1^; P=0.005; Figure 5B). The positive effect of trait correlations was consistent for all 6 traits; removal of trait correlations for each of the traits (by randomizing the values of one trait with respect to the others; see methods) produced similar distributions and significantly reduced performance compared to the observed combinations (Figure 7). Thus, the observed trait correlations are associated with higher feeding performance than the non-correlated case, but it is unlikely that they result from trait distribution along a performance ridge.

**Figure 7:**
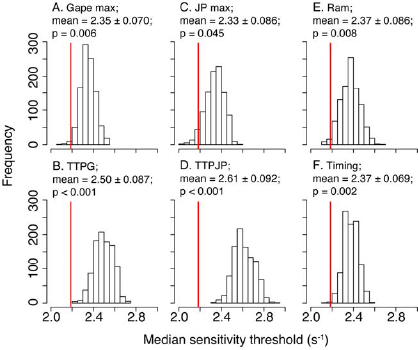
the effect of trait correlations on performance. We broke the correlation structure between the trait depicted in each panel and the other 5 traits and asked how the performance of these new combinations was affected. Mean denotes the average of median strain rate threshold for 1000 simulated populations; p value was calculated by comparing the distribution of median strain rate thresholds of the simulated populations with that of the observed one. Note that breaking the correlation for each single trait with the other five had similar effects across traits, reducing performance by 7.5-20%. Thus, the observed trait correlations are associated in higher feeding performance than the non-correlated case.

## Discussion

In this study, we refine the “performance landscape” framework suggested by Arnold (Arnold, 2003) by mapping the phenotypic distribution of performance-determining traits to a landscape derived from a theoretical model of a fitness-determining performance. Our approach allows a robust interpretation of the performance landscape beyond the ranges observed in the sampled population, and curtails “edge effects” that originally resulted in higher inaccuracy of the landscape near the population edge. Mapping the observed population on the landscape indicated that the population was located on the “upper slopes” of an adaptive peak, but not on the peak itself. Nevertheless, trait correlations did not seem to limit performance, as breaking trait correlations reduced performance. The major axes of phenotypic variation were tangent to the direction to the local performance peak, inconsistent with the scenario of trait distribution along a performance ridge (Fig 1). We conclude that our approach can help uncover the relationships between performance and trait evolution in complex functional systems.

### The Chromis viridis population is located off a performance peak

The performance landscape for suction feeding reported here was complex, rugged, and featured performance troughs and ridges (Figure 4). The landscape clearly showed that different phenotypic traits have a different effect on performance as indicated by the different slopes of the performance surfaces (Fig 4), and the different selection gradients for the different traits (Fig 6). Moreover, these slopes often change in relation to the values of the other traits. For example, for low values of time to peak jaw protrusion (TTPJP < 0.02 s), the effect of time to peak gape (TTPG) on performance is much stronger than at high values (TTPJP > 0.06 s; Fig 4). This is probably responsible for some of the variation in individual performance gradients (Fig 6). These trends demonstrate the complexity of the relationship between form and function in this functional system, and the importance of an integrative analysis to accurately model it.

Functional systems are often considered to be ‘optimized’ for performance, i.e. the distribution of phenotypic traits within a species or a population is assumed to correspond to performance peak (Bishop et al., 2008; van Leeuwen and Muller, 1984). This notion is often supported by using the measured performance data to construct a putative landscape (Arnegard et al., 2014; Arnold and Bennett, 1988). More often, and especially in an inter-specific context, it is the distribution of phenotypic traits themselves that is used to infer the existence and location of a “peak” (Collar et al., 2009; Ingram and Mahler, 2013; Shoval et al., 2012). However, the relationship’s correspondence (or lack thereof) between the performance landscape and the distribution of species traits are poorly demonstrated.

As pointed out by Arnold, (Arnold, 2003) it is problematic to infer the location of performance peaks from such data. Therefore, the location of the population with respect to performance peaks and ridges is difficult to resolve and is largely unclear. Here we show that suction feeding performance of *Chromis viridis* is not optimal, although it is significantly higher than that expected under a model of random motion across the landscape. Such a distribution is often explained by trade-offs with other performance axes. For example, Polly et al (Polly et al., 2016) argued that shell shape in turtles is determined by the trade-off between shell strength and form drag. However *C. viridis* is a zooplankton specialist, feeding mainly on copepods (Allen and Randall, 1980), and it is highly likely that the performance landscape can be reduced to one prey escape strategy (i.e. strain-sensitive prey). Unlike in mouth-brooding, nest building or biting/scraping fish species (Ostlund-Nilsson and Nilsson, 2004; Smith and Wootton, 1995), the mouth in *C. viridis* is used mainly for suction feeding and respiration. Therefore, it is unlikely that other fitness-determining functions impose strong direct selection on the traits we measured.

Instead, we speculate that suction feeding performance is constrained by mechanical limitations. Our model is purely functional, accounting for the effects of strike kinematics on performance. However, it does not account for energetic, genetic, developmental, or biomechanical constraints on the suction feeding mechanism (Carroll and Wainwright, 2009; Van Wassenbergh et al., 2006; Wainwright et al., 2007). Therefore, high-performance trait combinations identified by the model may be unrealistic due to such constraints. For example, biomechanical principles dictate that increases in gape size are likely to increase TTPG (Hulsey and Wainwright, 2002; Oufiero et al., 2012) and increase the energetic cost of the strike (Carroll and Wainwright, 2009). Such couplings are not accounted for by SIFF and are therefore not implicit in the landscape. Similarly, faster time to peak jaw protrusion might permit capturing more sensitive prey (Figure 4), however in reality it could be disfavored because it will increase the energetic cost of the strike (Carroll and Wainwright, 2009). Likewise, the relative timing of peak gape and peak jaw protrusion could be a consequence of the biomechanical lever system (Westneat, 2005), bearing little effect on performance within the observed range of trait values. Genetic linkages are also expected to constrain the distribution of traits in the observed population (Lande and Arnold, 1983; Schluter, 1996). In our simulated populations, we accounted for such correlations, which had a strong effect on the phenotypic distribution of our populations and on performance.

Alternatively, it could be that that the location of the population off the performance peak could result from a misalignment of the performance and fitness peaks. For example, it could be that the performance peak allows the capture of highly strain-sensitive prey. However, the energetic cost of feeding with a morphology that sits upon that peak is high. If the abundance of such highly evasive copepods is very low compared to other (less evasive) copepod species, the fitness peak (or peak energetic gain) will be shifted towards the current location of the population.

However, we are unaware of community-wide data on the distribution of hydrodynamic performance of copepods that would enable testing this idea.

The performance landscape is multidimensional, and it is therefore only possible to visualize it in two dimensional projections (Fig 4, Fig S2). Qualitatively examining the distribution of the observed population in these 2-D maps can be a useful tool in generating hypotheses regarding the distribution of the phenotypic traits on the landscape. For example, Figure 4D and 4F may support the intuition that some traits follow “performance ridges" (Arnold, 2003; Arnold et al., 2001; Ghalambor et al., 2003; Whibley et al., 2006), along which morphological variation can accumulate without performance consequences. However, it is important to remember that the 2¬D visualization only represents the behavior of the displayed variables at median (or another pre-defined) values of the other phenotypic traits, and cannot accurately represent trait distributions in the multidimensional space (Blonder, 2016; Gavrilets, 1997). The intuition that traits follow “performance ridges” is not supported by the individual selection gradients (Fig 6), which should have followed a hyperbolic shape (i.e. lower selection differential close to the population mean) if traits were located on a ridge. Additionally, the primary axis of variation was tangential to the path of steepest performance ascent, which is inconsistent with the “performance ridge“ hypothesis.

### Generality of the framework

Following the work of Hansen and Martins (Hansen and Martins, 1996), Arnold et al. (Arnold et al., 2001) suggested that using performance landscapes can be helpful in understanding micro and macro-evolutionary processes. However, such applications of the performance landscape were rare. For example, Tseng (Tseng, 2013) used finite element analysis to map two traits (skull width-to-length and depth-to-length ratios) to two functional properties, the mechanical advantage and strain energy of the Carnivora skull. That study calculated the strength of theoretical skull shapes to estimate the functional properties of traits that lay outside the distribution of observed skull shapes, concluding that skull shapes evolved towards higher performance, but also discovering high-performance regions that are not occupied by extant species. Similarly, Polly et al (Polly et al., 2016) used finite element analysis and hydrodynamic theory to predict the trade-off between shell strength and form drag, and mapped the distribution of turtle species with respect to the resulting Pareto front. By using the equations yielded from the landscape it is possible to relate mechanistic processes to divergence in trait values across intra- and interspecific levels: our results provide evidence for the evolution of feeding morphology on a complex performance landscape as an important process that drives the present intra-specific phenotypic distribution. It is possible to scale this method to higher phylogenetic levels, and test the role of the performance landscape in shaping inter-specific phenotypic distributions. Current methods for inferring adaptive regimes from morphological data use inter-specific trait variance to deduce the locations of “adaptive peaks" (Ingram and Mahler, 2013; Pennell and Harmon, 2013; Uyeda and Harmon, 2014). These methods do not require *a priori* assumptions regarding the manifestation of form to function, and the phylogenetic position, optimum, and relative “heights" of the adaptive peaks are inferred from the data. Using theoretical performance landscapes to detect adaptive peaks can be advantageous over such methods because (1) it can easily account for interactions between traits in multivariate space, (2) the accuracy of the landscape is independent of the sample size of the focal population, is unaffected by incomplete taxon sampling, and is uniform across the morphospace and (3) can estimate performance ridges and more complex nonlinear features of empirical landscapes, which cannot be generated by simple regime-shifting models such as the Ornstein-Uhlenbeck model (Hansen and Martins, 1996). Indeed, field experiments now demonstrate that complex fitness landscapes drive adaptive radiation in rapidly speciating fish groups (Arnegard et al., 2014; Martin and Wainwright, 2013). It will be interesting to compare the predictions of such methods with the topography of empirical performance landscapes.

Importantly, while our study focuses on suction feeding in fish, the framework we present here is general and widely applicable across taxa and functional systems. Over the last decades, biomechanical theory and computational methods were used to mechanistically model many aspects of performance such as swimming (Fish and Lauder, 2006; Liao, 2007; Sfakiotakis et al., 1999), running (Barasuol et al., 2013; Kingma et al., 1996; Minetti, 1998), slithering (Hu et al., 2009), flying and gliding (Paranjape et al., 2012; Wu, 2011) among numerous other examples. These models can be used to generate performance landscapes and map the distribution of functional traits on these landscapes in a variety of functional systems, and in many species. Because performance is fundamentally connected to viability selection in the wild, this framework now enables us to test how these complex functional systems shape evolutionary trajectories across species.

The performance landscape can be a useful tool to generate predictions regarding the functional consequences of shifting trait means in response to ecological (e.g. predation or competition) and evolutionary (e.g. co-evolution) processes (Arnold, 2003; Arnold et al., 2001; Ghalambor et al., 2003). For example, the performance landscape can predict how removing large individuals with larger gape (e.g. through predation or fishing) will change the distribution of performance in the population. Likewise, performance landscapes can be used to predict the morphological response to evolutionary changes in prey escape capabilities such as increased sensitivity or the ability to accelerate and swim faster. However, the complex nature of the landscape warrants caution when inferring selection, because selection may operate differently on the same traits in different areas of morphospace, depending on the values of other traits of the system. For example, two individuals that have the same trait value for maximal jaw protrusion can be subjected to different selection regimes, depending on the values of their five other traits. The selective force operating on each point in the multi-dimensional space can be calculated as the sum (for all traits) of the derivatives of performance with respect to changing trait values. It also follows that simple mathematical functions (Arnold et al., 2001) fail to capture the complex nonlinear features of the landscape (Schluter, 1996).

## Supplementary tables and figures

**Table S1:**
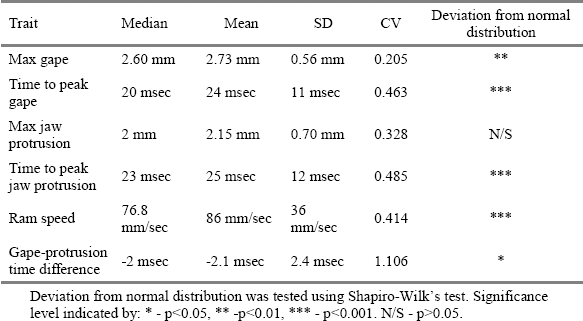
Summary statistics for the six morphological traits in the observed population

**Table S2:** Contribution of morphological traits and their interactions to the GAM- smoothed performance landscape.

| Trait | Effective degrees of freedom | Deviance from null (Wald statistic) | Significance level |
| --- | --- | --- | --- |
| Max gape (gape_max) | 5.10 | 211.625 | *** |
| Time to peak gape (TTPG) | 3.64 | 60.4 | *** |
| Max jaw protrusion (JP_max) | 5.82 | 153.039 | *** |
| Time to peak jaw protrusion (TTPJP) | 6.72 | 74.413 | *** |
| Ram speed (ram_spd) | 5.58 | 2596.523 | *** |
| Gape-protrusion time difference (time_diff_peak) | 4.02 | 7.095 | *** |
| gape_max:TTPG | 2.48 | 4.854 | ** |
| gape_max:JP_max | 7.17 | 3.981 | *** |
| gape_max:TTPJP | 10.88 | 6.721 | *** |
| gape_max:ram_spd | 10.53 | 45.606 | *** |
| gape_max:time_diff_peak | 5.68 | 2.235 | * |
| TTPG:JP_max | 0.00 | 0.481 | N/S |
| TTPG:TTPJP | 11.47 | 23.131 | *** |
| TTPG:ram_spd | 7.39 | 18.293 | *** |
| TTPG:time_diff_peak | 13.44 | 12.051 | *** |
| JP_max:TTPJP | 10.74 | 12.511 | *** |
| JP_max:ram_spd | 7.49 | 33.797 | *** |
| JP_max:time_diff_peak | 3.27 | 1.573 | N/S |
| TTPJP:ram_spd | 12.22 | 14.315 | *** |
| TTPJP:time_diff_peak | 11.58 | 34.186 | *** |
| ram_spd:time_diff_peak | 9.20 | 2.318 | ** |

3 Wald statistics and p-values were calculated according to Wood (2013). Significance 4 level indicated by: \* - $p < 0.05$ , \*\* - $p < 0.01$ , \*\*\* - $p < 0.001$ . N/S - $p > 0.05$ .

**Fig S1:**
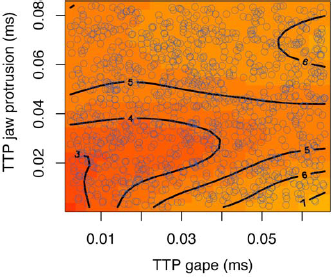
The distribution of 3291 phenotypic combinations in a 2D data projection (time to peak gape and time to peak jaw protrusion). Trait combinations (blue circles) were generated by randomly sampling from the range of phenotypic traits observed in the population, and used to generate the landscape (black contours and red-yellow colors).

**Figure S2:**
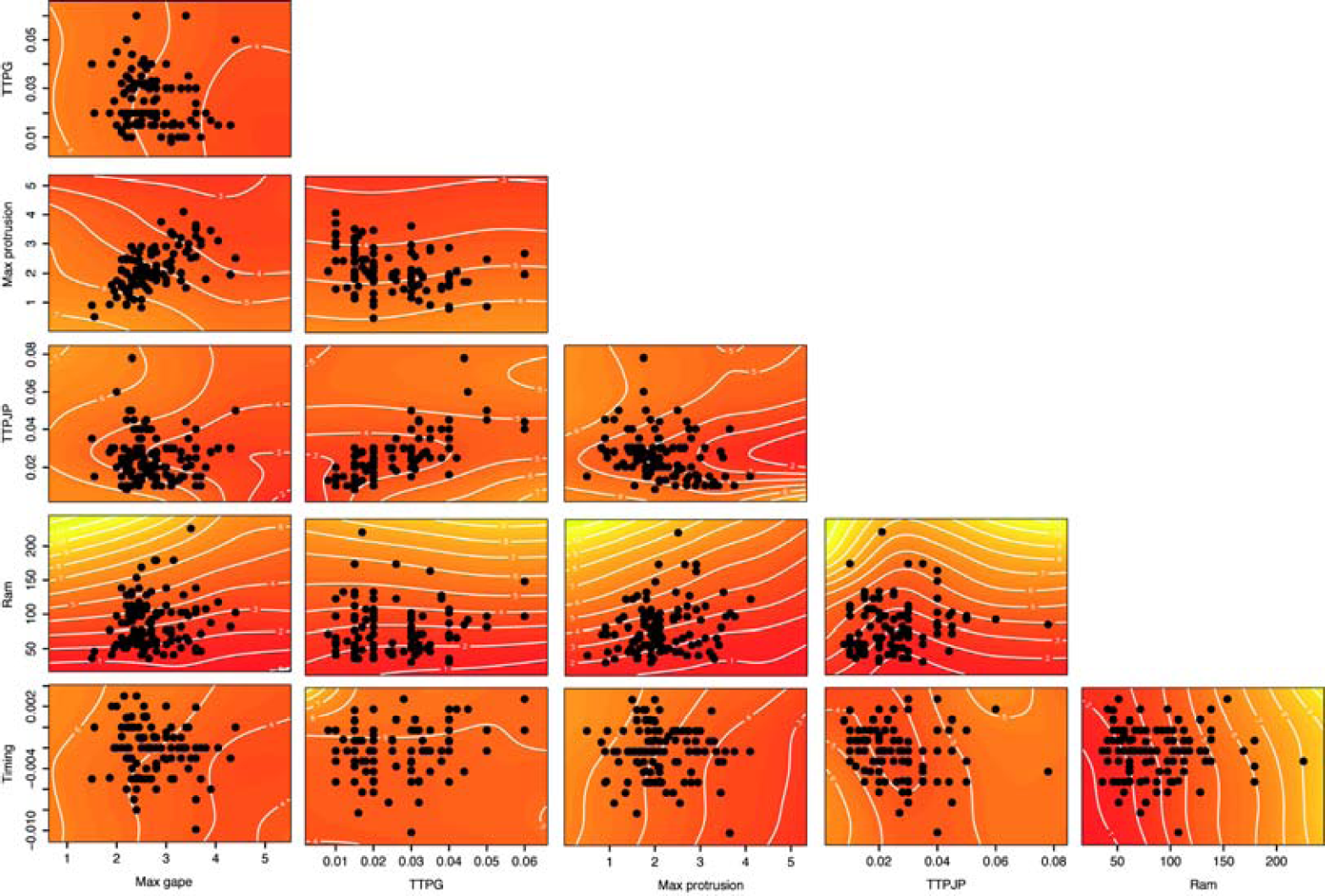
2D projections of the multidimensional performance landscape, generated using a hydrodynamic model of predator-prey interactions. The plot features all possible 2D projections available for the six traits measured. The multidimensional performance landscape was generated based on 3300 simulated prey-acquisition strikes, featuring random trait combinations selected from the observed trait range. Colors (and white contours) represent feeding performance, estimated as strain rate threshold (s^−1^) of the most sensitive prey that can be captured by a predator using that trait combination. Red colors represent high performance (low strain rate thresholds), yellow represents low performance (high strain rate thresholds). Observed trait combinations (black circles, n=110) are overlaid on the performance surface. Note that the 2D visualizations only represent the performance of the featured variables at median values of the other four phenotypic traits, and do not represent trait distribution in the multidimensional space.

